# Hip retraction enhances walking stability on a ramp: an equilibrium point hypothesis-based study

**DOI:** 10.1101/638635

**Authors:** Alireza Bahramian, Elham Shamsi, Farzad Towhidkhah, Sajad Jafari

## Abstract

Hip retraction is a phenomenon observed in human walking. The swing leg rotates backward at the end of the motion. Its positive effect on motion stability was reported in the literature based on some simple models for running or walking. In this study, it is shown that hip retraction angle increases in humans during their ascending and descending walk on a stair. In previous studies, hip retraction was modeled by defining a proper motion for the swing leg. According to the equilibrium point hypothesis, the central nervous system (CNS) defines only the equilibrium point(s) and stiffness(es) for body joint(s) to control the human motion. Human body motion emerges as its natural response as a result of the external forces and the defined equilibrium points of joints. Considering the hip torque as a spring-like model with an equilibrium point and stiffness, this study revealed that the hip retraction can be generated by the natural response of the swing leg. Besides, the stabilizing effect of hip retraction was demonstrated by a model for human’s ascending and descending walking on a ramp with a range of positive and negative angles, respectively. The findings suggest that the CNS needs to define equilibrium point just ahead of the stance leg to take advantage of the hip retraction effect on ascending and descending walks on a ramp.

## 1. Introduction

Human leg retracts at the end of the swing phase just before the leg collides with ground (i.e., heel-strike) [1]. It was shown that hip retraction exists in human motion for a wide range of walking and running speeds [2]. Leg retraction was also observed in animal locomotion [3]. Several positive effects, such as motion stabilization and energy optimization, were mentioned for this phenomenon [4].

Motion stability is one of the important issues that has been studied from different viewpoints [5–7]. Positive influence of hip retraction on the motion stabilization was demonstrated via mathematical models [8] and robots [9]. Fixed retraction angular velocity for swing leg which planned to begin at the mid-flight, increased the running model stability [10]. In addition, using a predefined retraction angle for the swing leg improved the stability of the simple inverted pendulum model for a passive walking on downhill [11]. In another study, the effect of hip retraction on the stability of a walking robot disturbed by a stepdown was investigated [12]. This study also showed that using a predefined mild retraction trajectory for the swing leg resulted in an optimal disturbance rejection.

Optimality is another important factor in locomotion that has frequently been investigated in literature [13, 14]. Another major benefit of hip retraction is reducing energy consumption. It helps to reduce energy loss by declining the relative speed between the ground and the swing leg [15]. This was also confirmed by some model-based studies. For example, modeling hip retraction with an impulse that moved the swing leg downward, helped to decline energy losses at heel-strike during a high-speed walking [16]. In another study, optimal timing of hip retraction to increase energy efficiency was investigated by a similar model [17, 18].

Moreover, the retraction rate during running was optimized for making use of both stabilizing and enhancing energy efficiency of hip retraction [19]. Improving state estimation was mentioned as another benefit of hip retraction [20].

Introduced by Feldman, Equilibrium Point Hypothesis (EPH) is one of the well-known and impressive theories in the field of motor control. In this theory, it is assumed that the Central Nervous System (CNS) sets the reference muscle length (λ) for each muscle [21, 22]. According to EPH theory, muscle force is not programmed. It emerges on the basis of the difference between the real muscle length and the muscle referent, instead. This theory was then developed into the body referent theory [23, 24]. In the body referent theory, two main parameters (i.e., referent and stiffness) are defined for each joint [25]. CNS only determines these parameters instead of programming a certain motion. The motion is emerged based on the defined parameters and external forces. According to the theory, it is assumed that a mapping from some global parameters builds the referent and stiffness for each joint. Global referent (i.e., body referent location) and global stiffness are suggested as global parameters for a multi-joint system [26]. Existence of such global parameters is necessary to provide an integrated motion and coherent muscles. Global electromyogram (EMG) minima is an evidence for the existence of such global parameters. It occurs at the moment that most of the muscles are at their lowest activation state rather than their maximum magnitude [27]. This phenomenon was observed in several motor tasks for humans [28] and animals [29]. Global EMG minima is interpreted at the moment that the global referent collides with the real body position. This is because most of the muscles are close to their references when they are at their lowest activation state. A global EMG minima was reported for stepping forward at the moment that the swing hip is going a little ahead of the stance hip [30].

In this study, hip retraction for human stepping on flat ground, ascending and descending ramps are investigated. A double pendulum model is used for modeling the walking. Two controllers for fixing step length and duration are needed for comparing walking stability on a ramp, with and without hip retraction. First, the effects of the model parameters on the step length and duration are investigated and analyzed. Then, the controllers for the model are designed considering the results of analysis. Dependence of the hip retraction on the assumed equilibrium point for hip is shown afterwards. Finally, hip retraction effects on the walking stability on positive and negative ramps (i.e., ascending and descending walk, respectively) are evaluated.

## 2. Material and Methods

### 2.1. Experimental data

A dataset of a relevant research [31] was used to investigate hip retraction during the stepping on flat ground, ascending, and descending stairs. The data were recorded from 20 healthy subjects (age: 6-17 years, mean age: 10.8 ± 3.2 years; body mass: 41.4 ± 15.5 kg; height: 1.47 ± 0.20 m; 9 males, 11 females; for more details about the dataset, see [31]). Data of the normalized stride on the flat floor is illustrated in Fig.1. It can be seen that the hip flexion angle increased to 35.5^°^ and then it experienced a decrease of 3.6^°^.

**Fig. 1.**
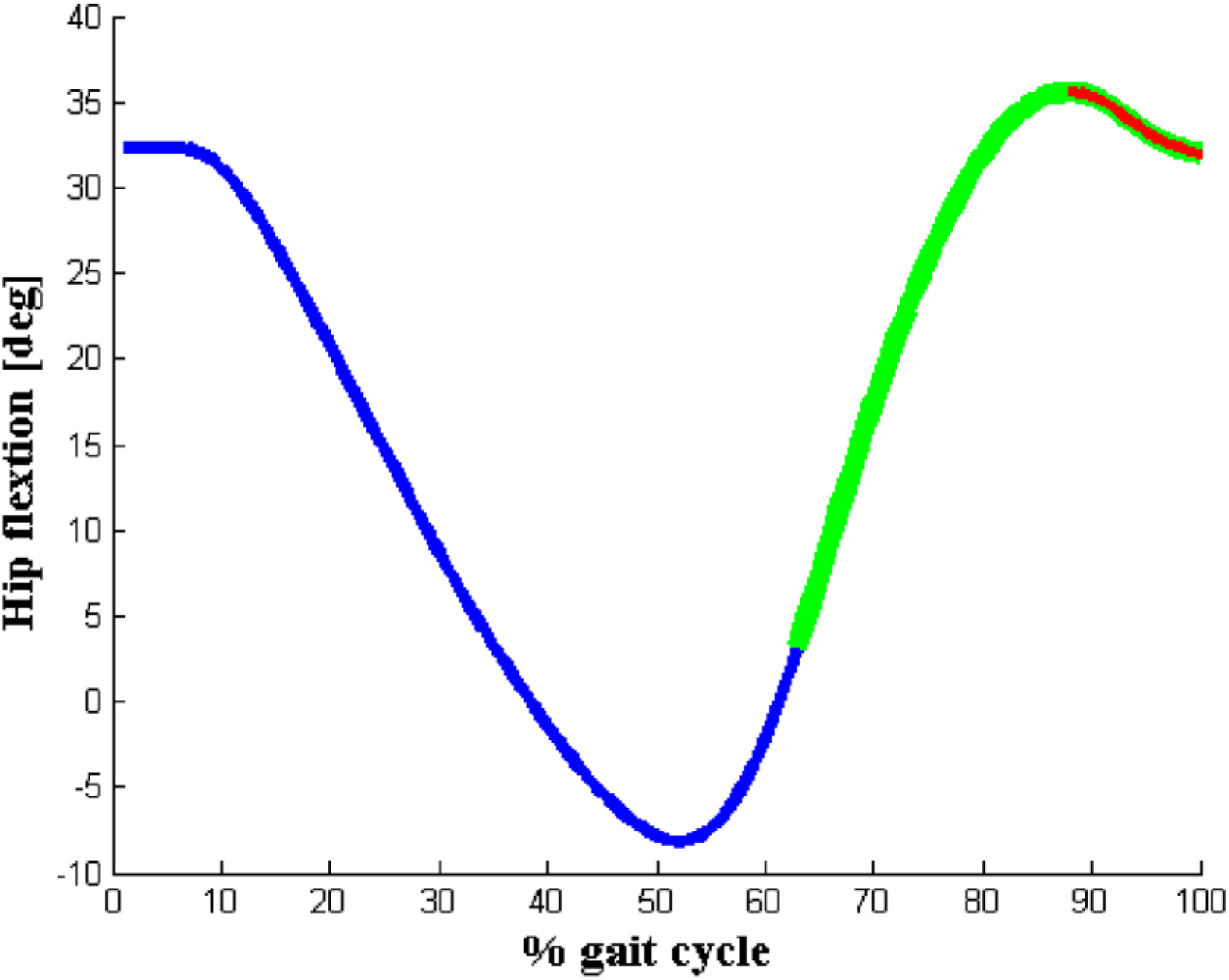
Hip flexion for the normalized stride on flat ground (i.e., heel-strike to heel-strike) (blue: stance phase, green: swing phase, red: hip retraction at the end of the swing phase). Retraction angle was about 3.6^°^.

Normalized hip flexion data for stride on an ascending stair is displayed in Fig.2. In this state, hip flexion angle went up to 38.8^°^ and the retraction of the swing leg was 11.0^°^. As it is shown, value of hip retraction for the ascending stair compared to stride on the flat floor increased.

**Fig. 2.**
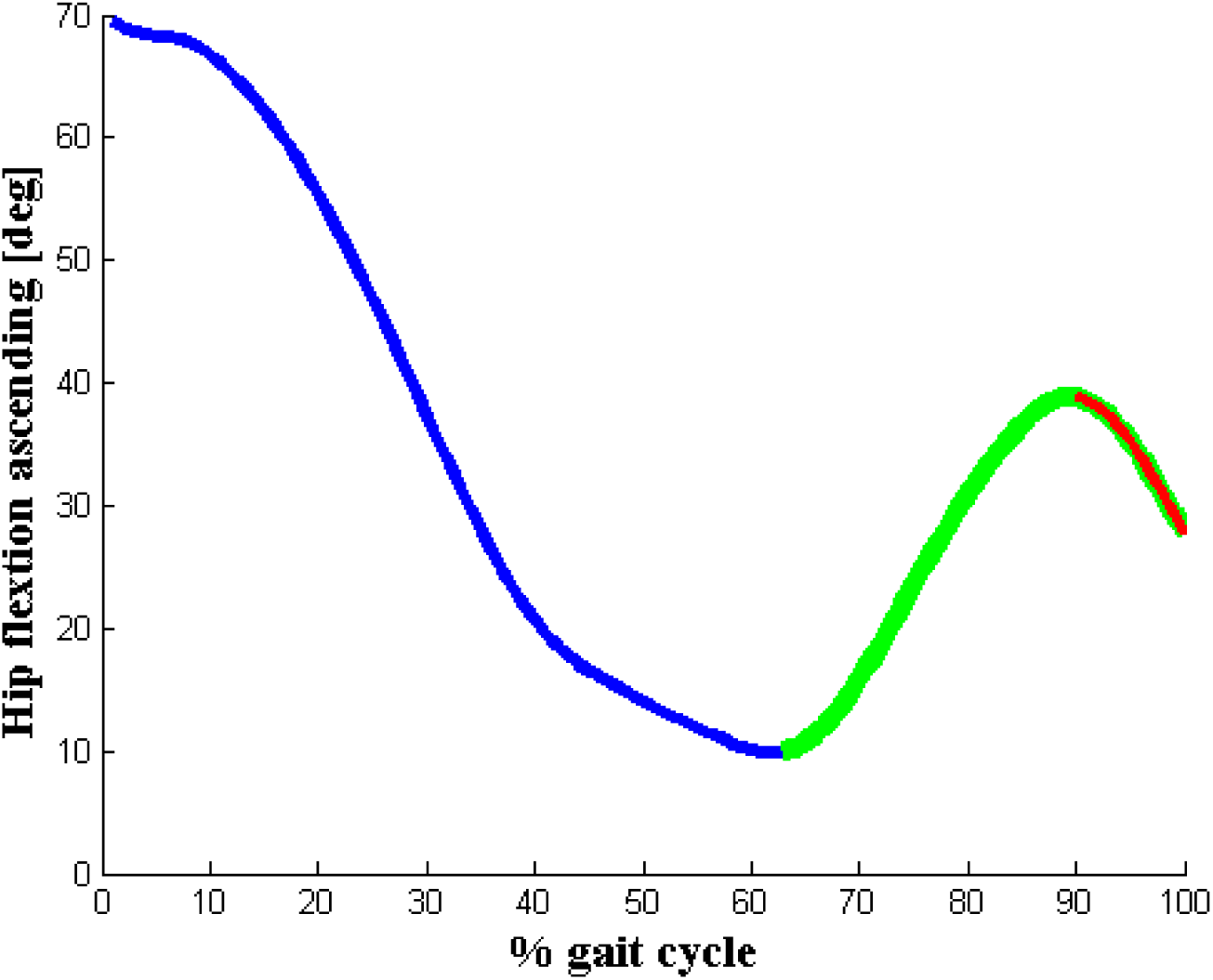
Hip flexion for the normalized stride on an ascending stair (blue: stance phase, green: swing phase, red: hip retraction at the end of the swing phase). Retraction angle was 11^°^.

Normalized hip flexion data for the stride on a descending stair is displayed in Fig.3. The figure shows that the hip angle first rose to 40.4^°^, then to 11.2^°^ retraction. The hip retraction was higher than that of the flat floor for the ascending and descending stairs.

**Fig. 3.**
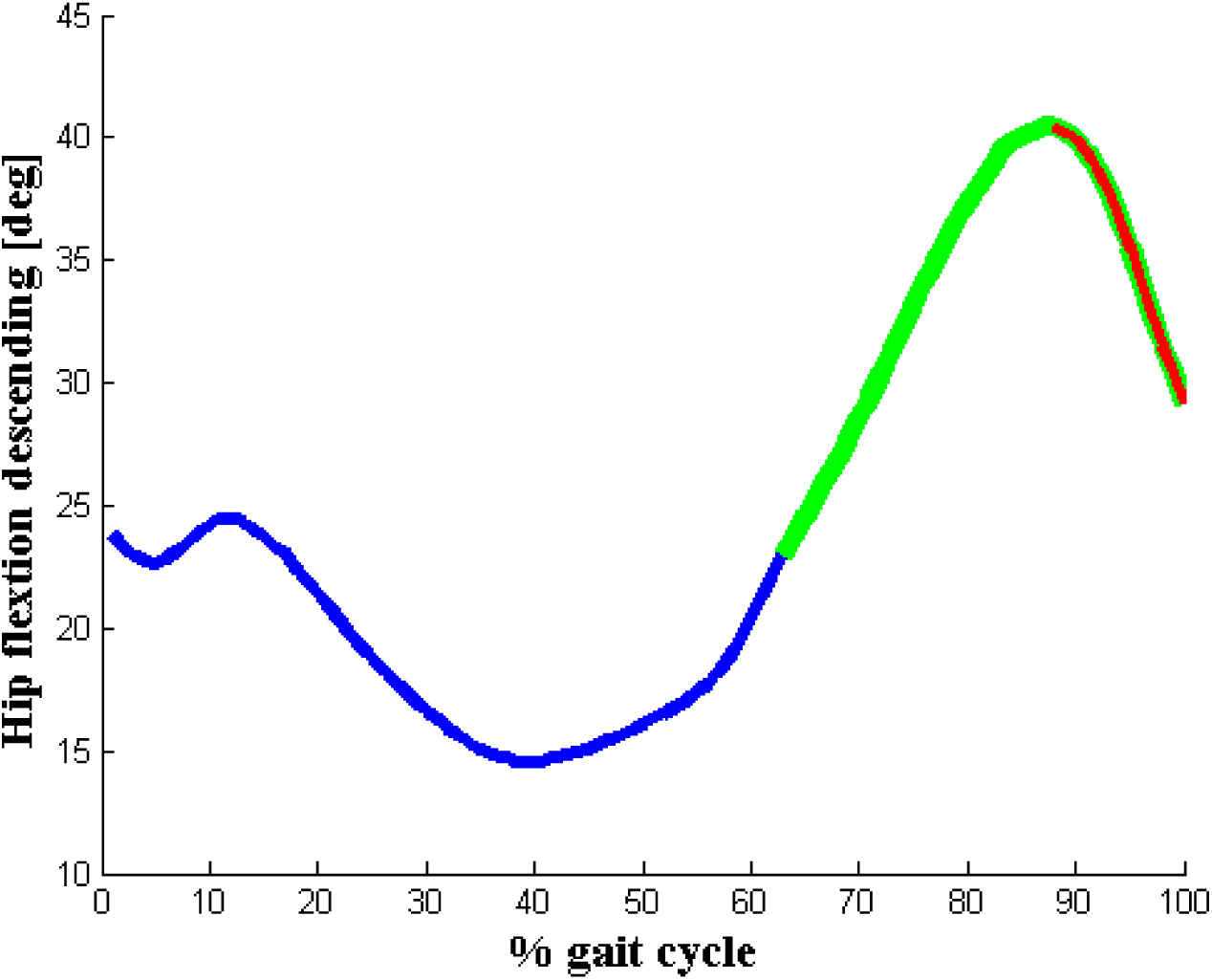
Hip flexion for the normalized stride on a descending stair (blue: stance phase, green: swing phase, red: hip retraction at the end of the swing phase). Retraction angle was 11.2^°^.

It seems that the hip retraction enhances the stability of walking on ascending or descending stairs. Since a large number of studies focusing on hip retraction have used simple conceptual models, a common model was utilized for present study, as well. We analyzed how the hip retraction influences the walking stability on the ramp, which seems to have some similarities with the stride on the stair.

### 2.2. The proposed model

A common double pendulum was used as a model of biped and *l*=1 was considered as the leg length. Dimensionless variables were utilized with the following basic units: overall mass *M*, and the gravitational constant *g*. Then, time was normalized by 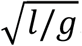. Eq.(1) and Eq.(2), which were frequently used in the literature [32–34], formed the model for simple biped walking based on the aforementioned scaling. The equations modeled the single support phase of walking. They were calculated as follows:

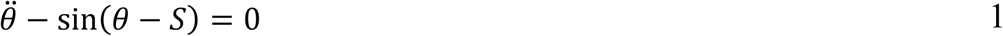

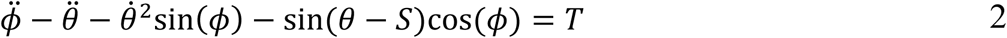

where *S*, *θ*, and *ϕ* were the ramp angle, stance leg angle, and swing leg angle, respectively. Besides, *T* was torque of the swing hip, which was considered as a spring with an equilibrium point (*ϕ*_0_) and a stiffness (*k*) (Eq.(3)).

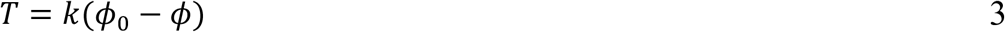

The condition for the end of the single support phase is mentioned in Eq.(4). The swing leg collides with the ground when the condition is met [34]

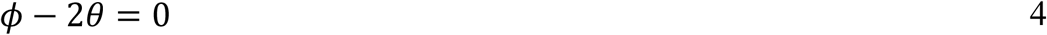

Additionally, Eq.(5) may be considered as the double support phase in which the swing leg and the stance leg are switched alternately. Collision with the ground affects the leg velocity. The push-off impact (*Impulse*) also influences the velocity of each leg. It should also be noted that the superscript + in Eq.(5) refers to the initial state of the variables to start the next step, whereas the superscript – is the final state of the previous step.

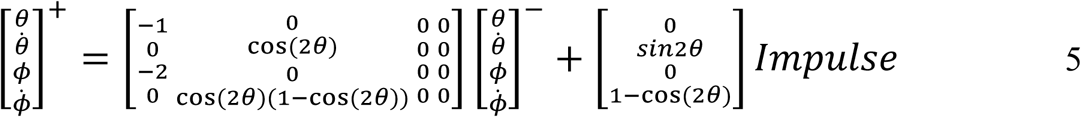

A schematic representation of the single support (i.e., Eq.(1) and Eq.(2)) and double support (i.e., Eq.(5)) phases are displayed in Fig.4.A and Fig.4.B, respectively.

**Fig. 4.**
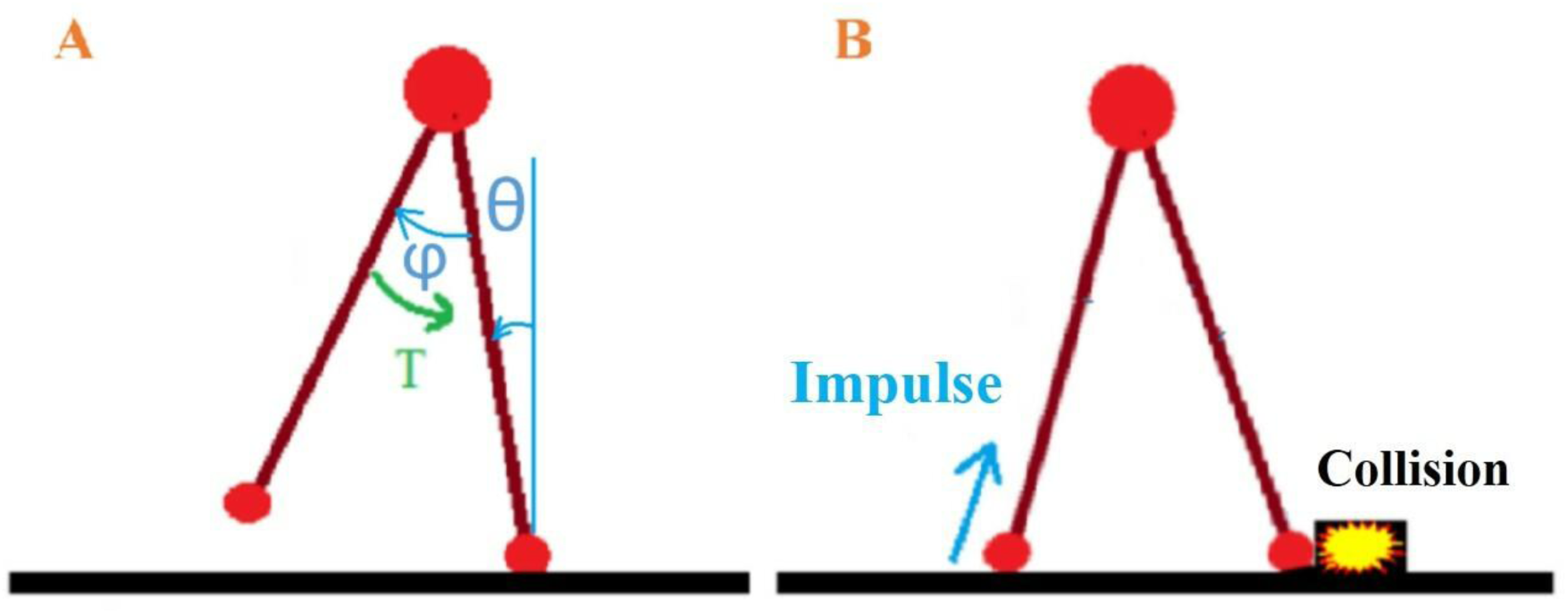
Schematic representation of single and double support phases. A) Single support phase (In this phase, torque is applied to hip from the assumptive torso). B) Double support phase (*Impulse* is applied to the model in this phase. In addition, collision with the ground and alternate switching between legs were considered in Eq.(5) for this phase).

So far, three parameters (i.e., *k*, *ϕ*_0_, and *Impulse*) were existed in the considered model. Fig. 5 illustrates the steady state value of *θ* and *ϕ* (*k* = 40, *ϕ*_0_ = −0.02, *Impulse* = 0.3). Step duration and length were calculated at the end of each single support phase.

**Fig. 5.**
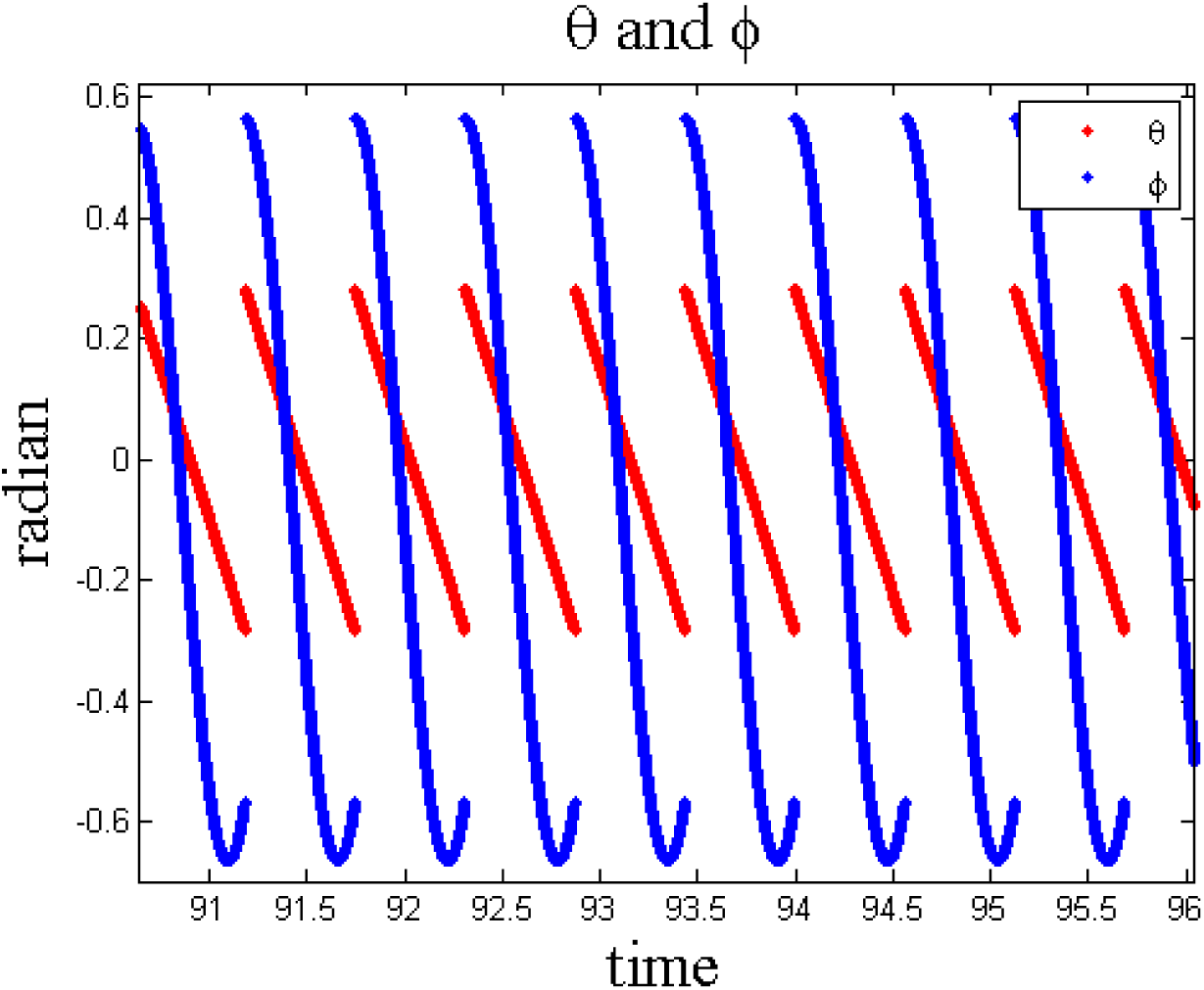
Steady state angles (blue: swing leg (*ϕ*), red: stance leg (*θ*)).

To design the step length and duration controllers for walking, it is necessary to identify effects of the model parameters on them. For this purpose, first, one parameter was considered fixed. Then, the values of two other parameters were set. The value of the step duration and length were computed after the transition mode. The effects of both parameters on step length and duration were investigated by setting a range of values.

Considering this approach, the equilibrium point of the swing hip (*ϕ*_0_) was fixed at −0.02. Then, two other parameters (*Impulse* and *k*) were set at different values. Step duration of model for different values of the parameters is shown in Fig.6. The effect of *k* on the step duration was much more than that of *Impulse*. The values of the step length for the same range of the two parameters are shown in Fig.7. It can be inferred that step length was much more affected by *Impulse* than *k*.

**Fig. 6.**
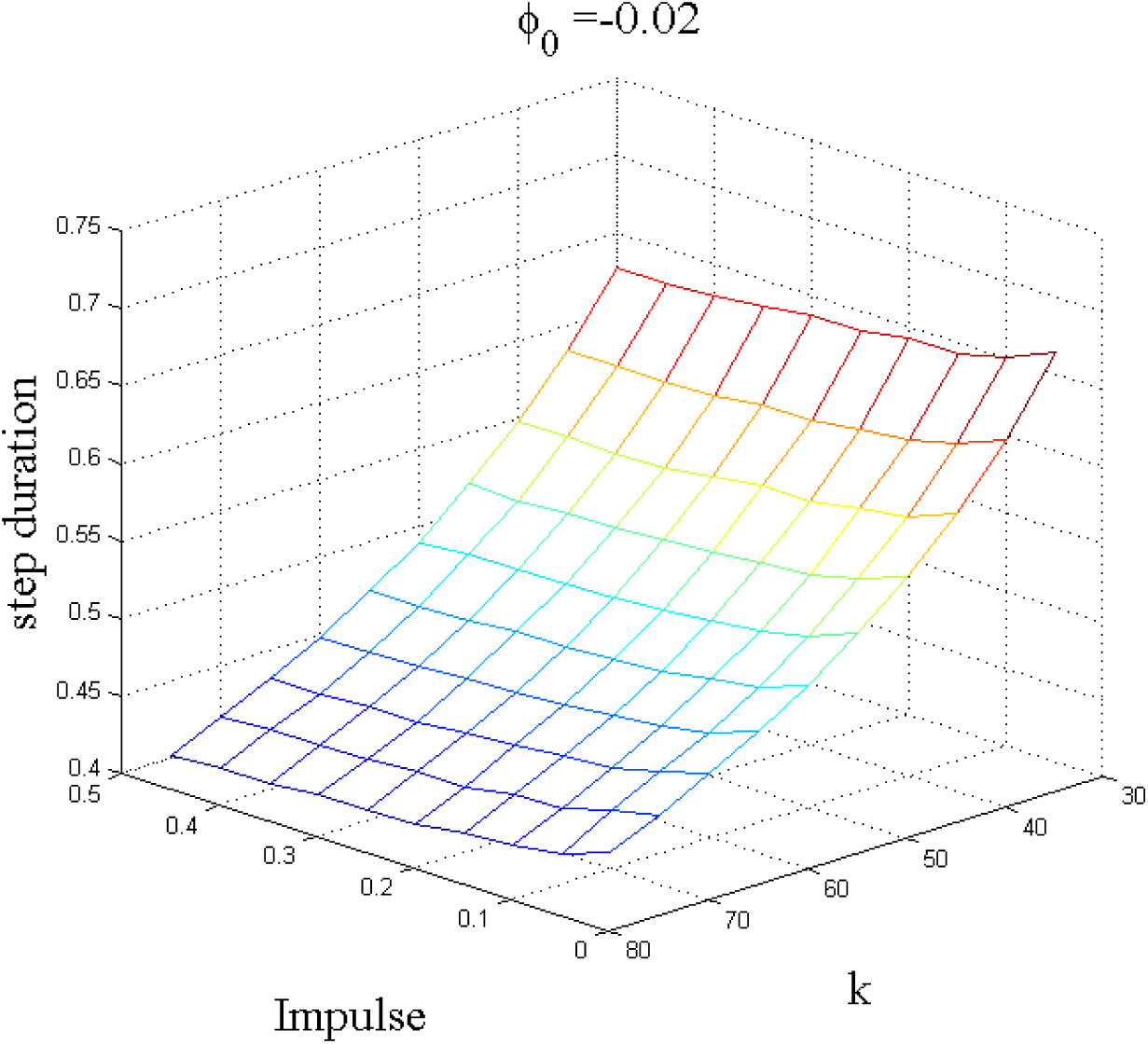
Step duration for different values of *Impulse* and *k* (*ϕ*_0_ was fixed at −0.02). Step duration was mainly influenced by *k*.

**Fig. 7.**
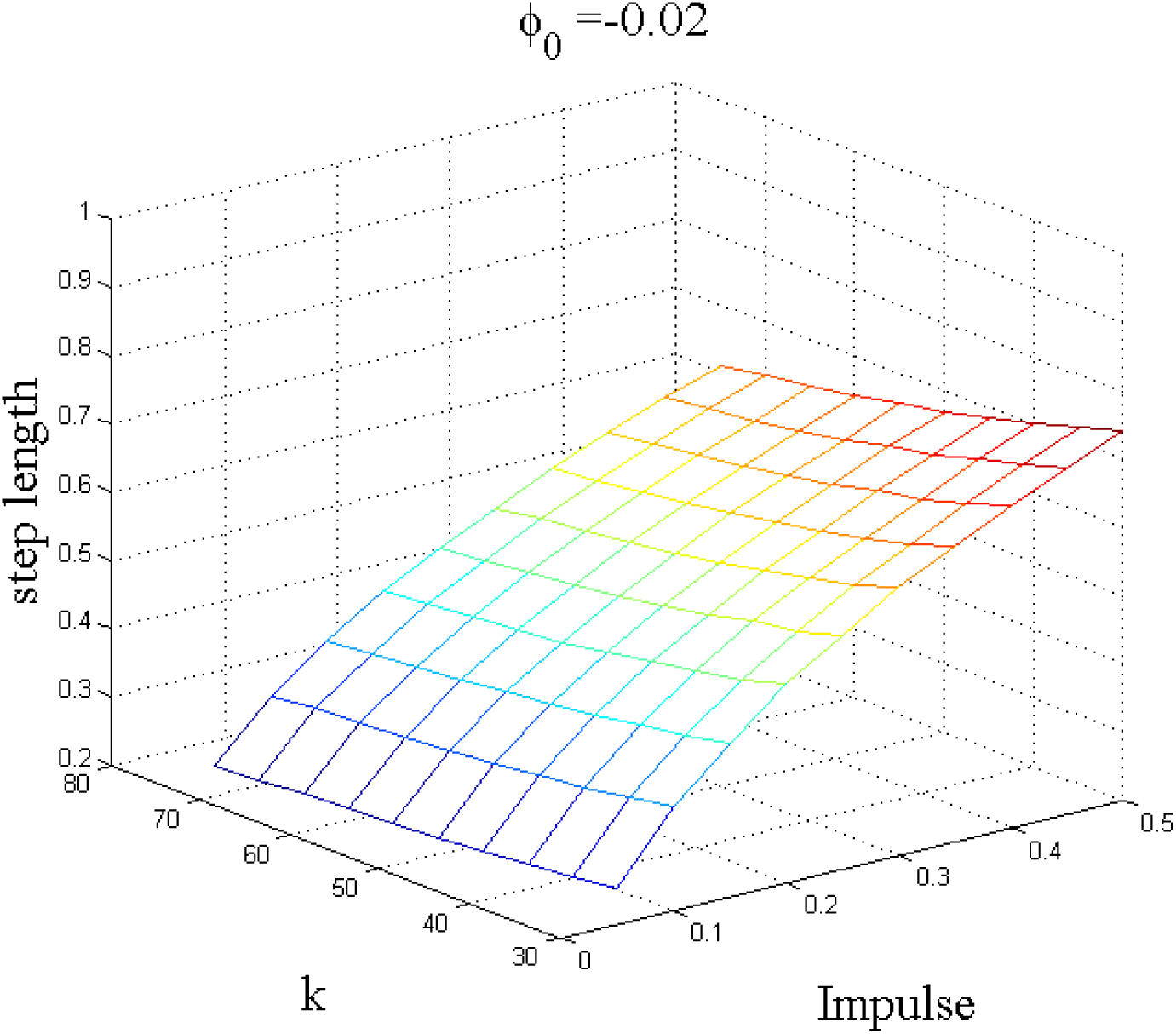
Step length for different values of *Impulse* and *k* (*ϕ*_0_ was fixed at −0.02). Step length was mainly influenced by *Impulse*.

### 2.3. Equilibrium point and swing leg retraction

Degree of the Swing Leg Retraction (SLR) was defined as the degree of retraction of *ϕ* after the swing leg arrived at its farthest front position (i.e., minimum value of *ϕ*). Degree of SLR was calculated by Eq.(6). Fig.8 displays the degree of SLR and the absolute value of *ϕ* in its farthest front position (i.e., maximum magnitude of *ϕ*).

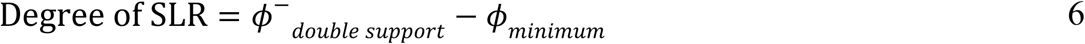

**Fig. 8.**
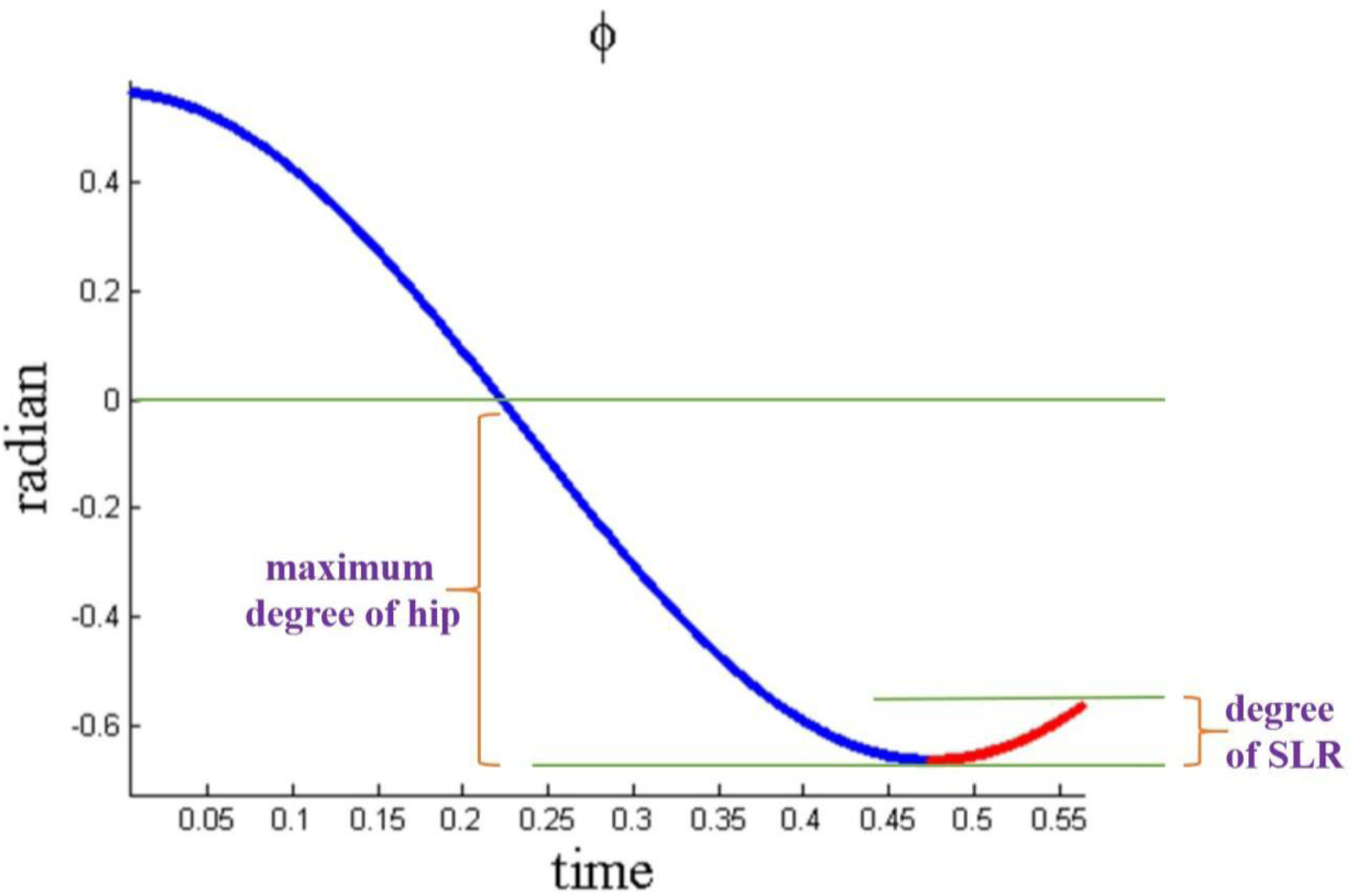
Swing leg angle (*ϕ*) (After the swing leg arrives at its farthest front position, it moves backward (i.e., retracts). Retraction is in red).

Initially, one parameter was considered fixed at its steady state value to investigate how the parameters of model affect the degree of SLR and maximum degree of hip. Then, values of the other parameters were set. The value of SLR and maximum degree of hip were calculated after the transition mode. The effects of parameters on the SLR and maximum degree of hip were investigated by setting a range of values for them.

### 2.4 Designing step length and step duration controllers for the model

Controllers were designed for the step length and duration in order to compare walking stability on a ramp with and without hip retraction. As it is previously shown, the step duration and length were mainly influenced by *k* and *Impulse*, repectively. Block diagram of the controllers for the step duration and length is illustrated in Fig.9.

**Fig. 9.**
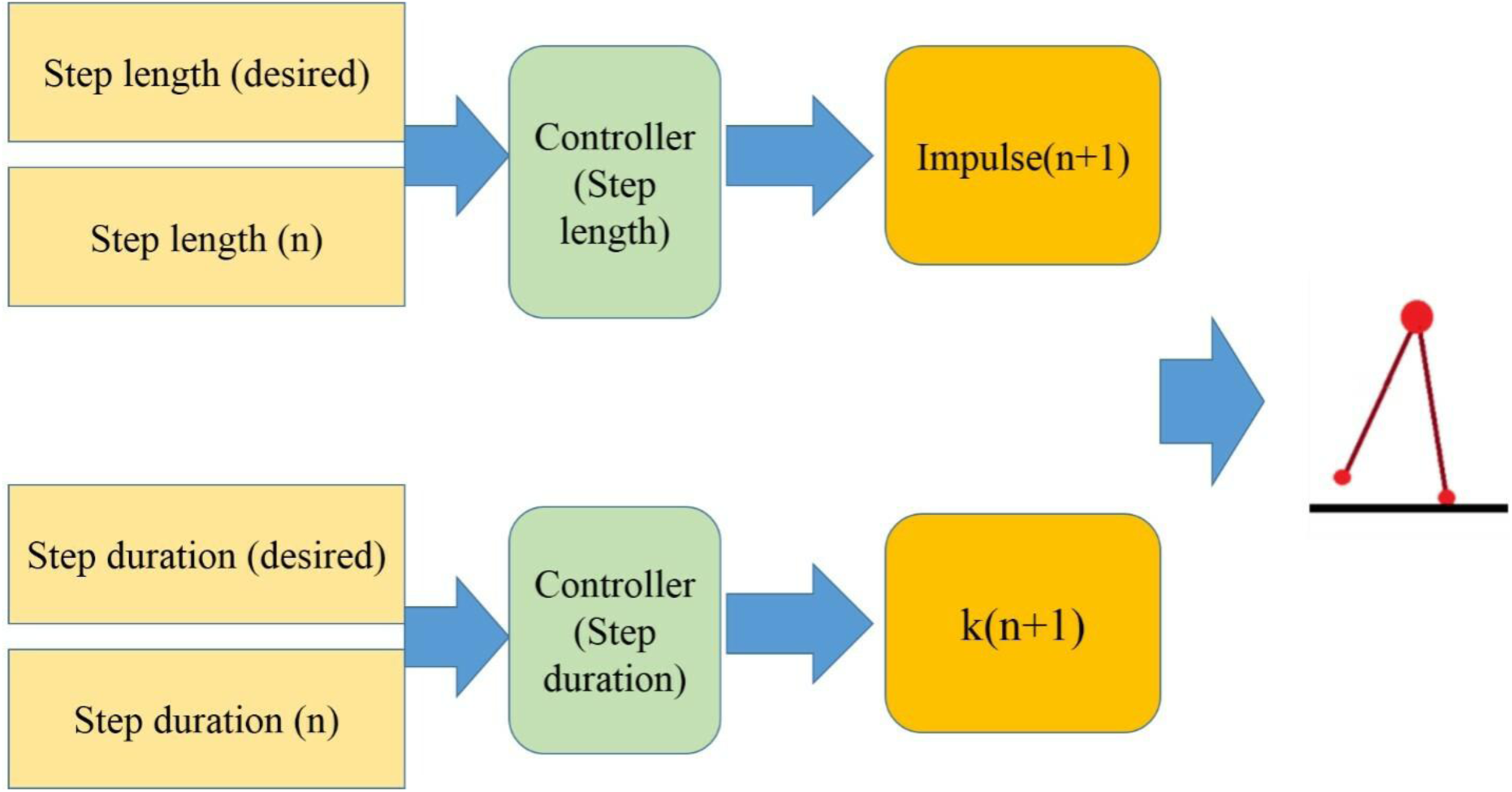
Block diagram of the controllers for the step length and the step duration. At the end of each step in the double support phase, based on the comparison between the last step length and the desired step length, *Impulse* for the next step is determined. Similarly, *k* for the next step changes on the basis of the comparison between the last step duration and the desired step duration.

Afterwards, an event-based controller was designed for the step length to reach its proper value according to Eq.(7) and Eq.(8). The last step length and the desired step length were compared in Eq.(7). Based on this comparison, *Impulse* for the next step was changed. The same structure of controller was used in a relevant study, namely *neural signal controller* [35].

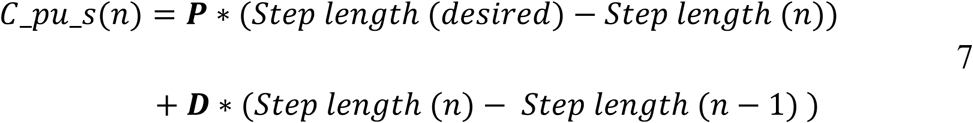

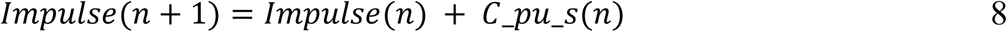

Likewise, *k* for the next step duration was determined (Eq.(10)) according to the comparison between the last step duration and the desired step duration (Eq.(9)).

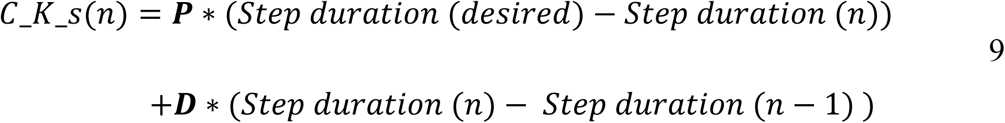

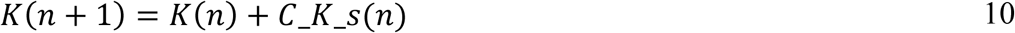

## 3. Results

### 3.1. Dependence of the SLR degree and maximum degree of hip on the model parameters

*Impulse* was fixed at 0.3 and then the other parameters (*k* and *ϕ*_0_) were set at different values. Fig.10 shows the effect of these parameters on the degree of SLR. It can be seen that the degree of SLR was mainly under the influence of *ϕ*_0_. Moreover, the maximum degree of hip increased with increasing the magnitude of *ϕ*_0_. Besides, decreasing the value of *k* led to the growth of the maximum degree of hip (Fig.11).

**Fig. 10.**
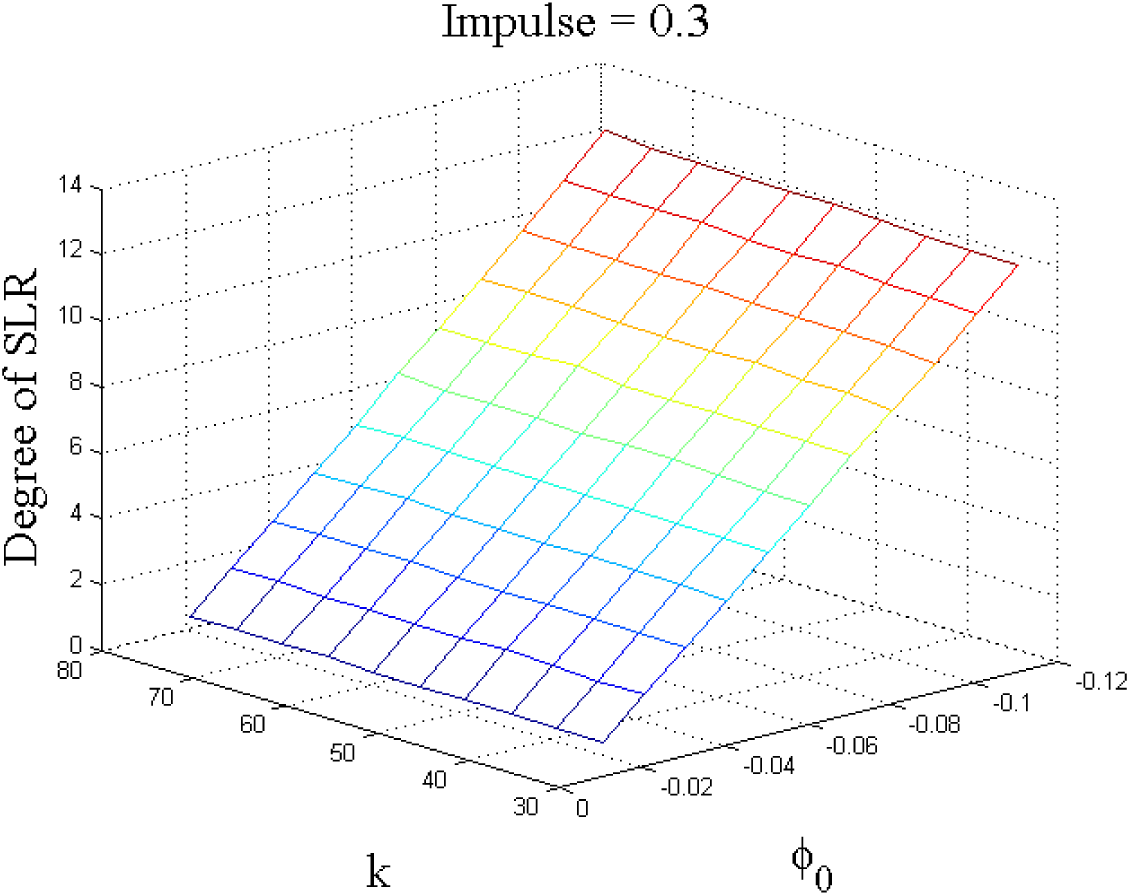
Degree of SLR for different values of *ϕ*_0_ and *k* (The *Impulse* was fixed at 0.3). As it can be seen, the degree of SLR was mainly affected by *ϕ*_0_.

**Fig. 11.**
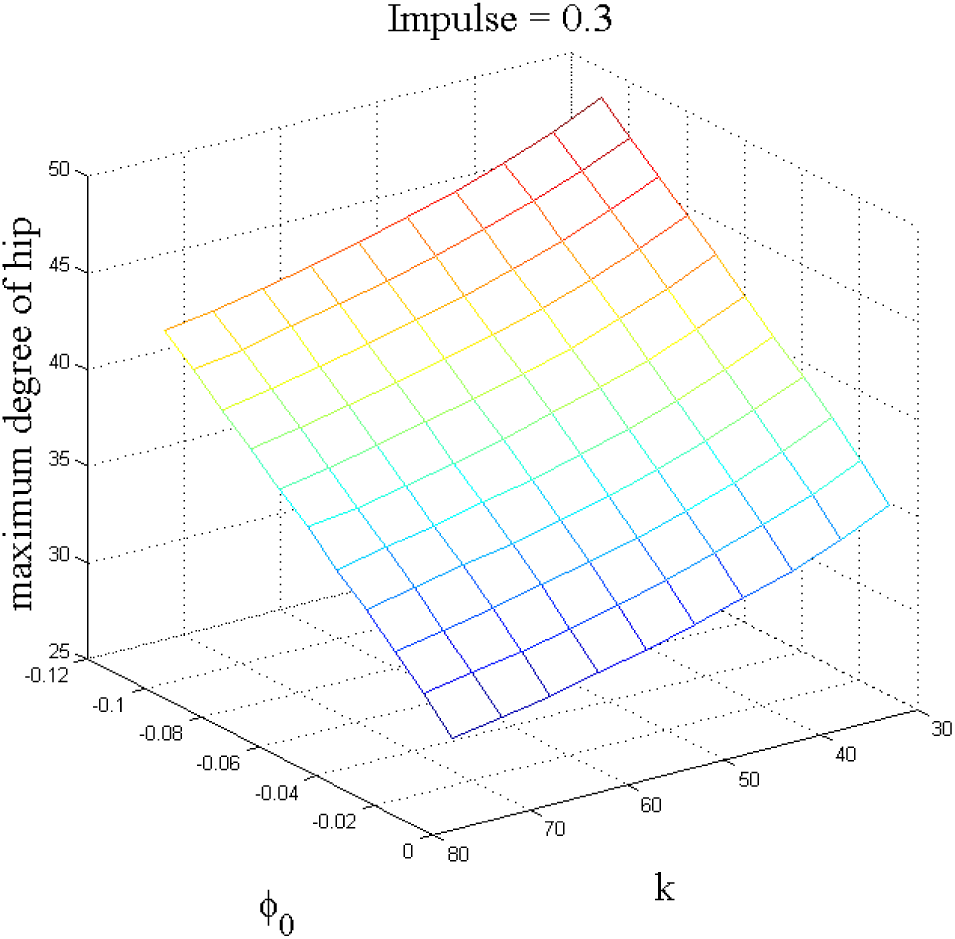
Maximum degree of hip for different values of *ϕ*_0_ and *k* (*Impulse* was fixed at 0.3). Decreasing *k* and increasing the magnitude of *ϕ*_0_ enhanced the maximum degree of hip.

Likewise, *k* was fixed at 40, and the degree of SLR was computed by setting different values for *ϕ*_0_ and *Impulse* (Fig.12). Similarly, the degree of SLR was chiefly influenced by *ϕ*_0_. Maximum degree of hip climbed with rising both the magnitude of *ϕ*_0_ and *Impulse* (Fig.13).

**Fig. 12.**
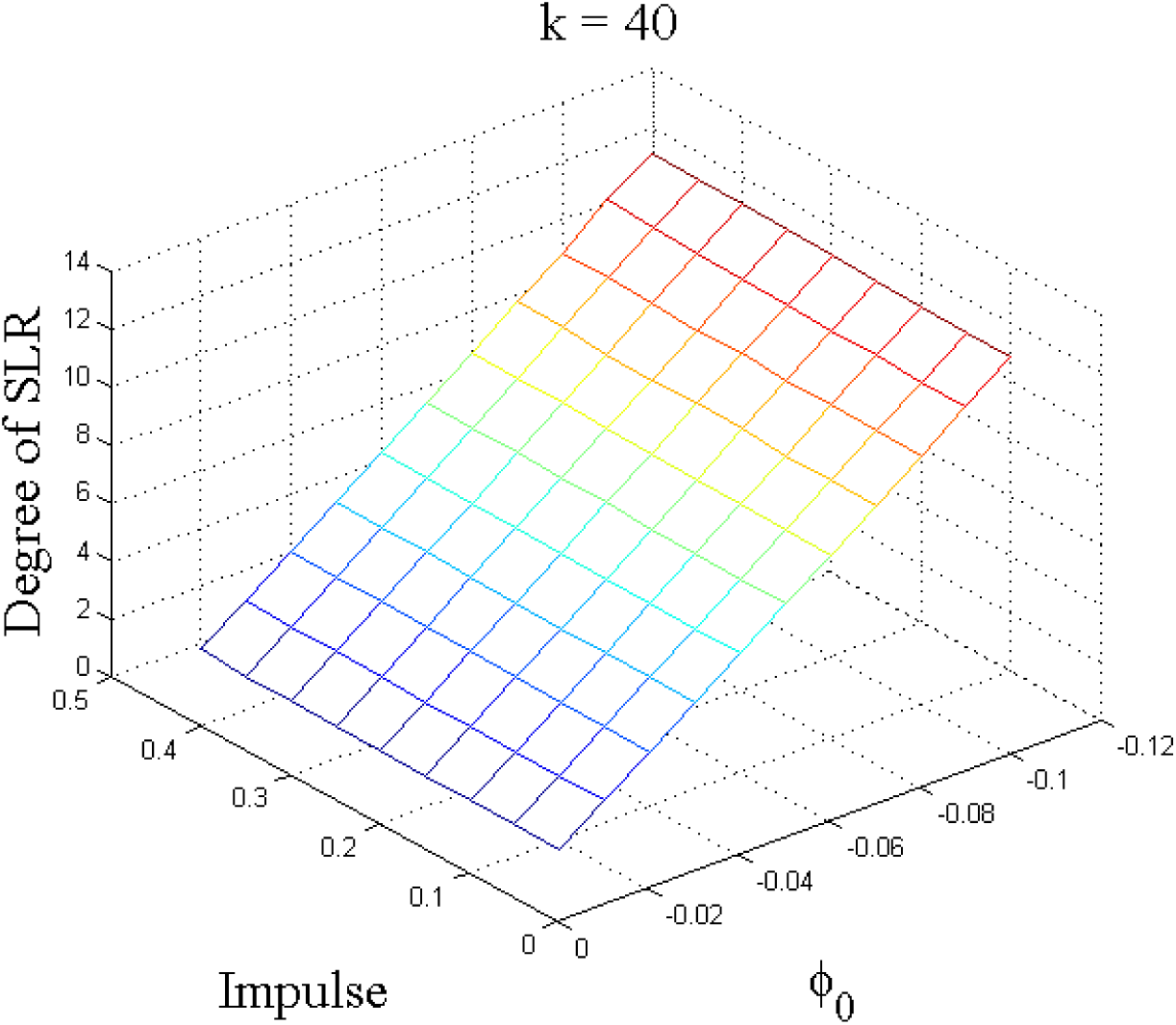
Degree of SLR for different values of *ϕ*_0_ and *Impulse* (*k* was fixed at 40). As it can be seen, the degree of SLR was mostly affected by *ϕ*_0_.

**Fig. 13.**
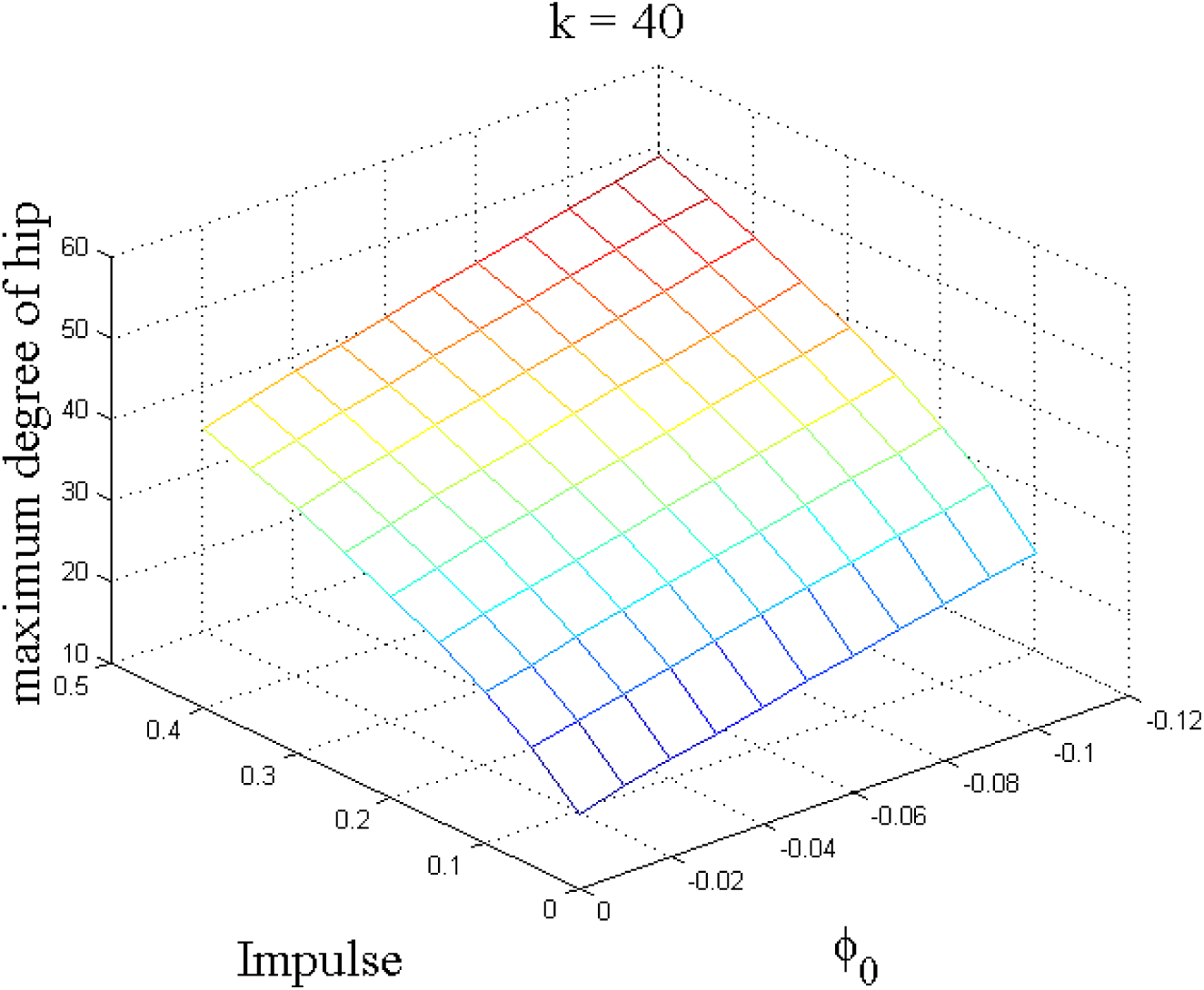
Maximum degree of hip for different values of *ϕ*_0_ and *Impulse* (*k* was fixed at 40). Rising both *Impulse* and the *ϕ*_0_ magnitude increased the maximum degree of hip.

### 3.2. Walking on the ascending and descending ramps

Table 1 shows the parameters of the designed controllers, which were configured manually. The desired step length was set at 0.5 and the desired step duration was adjusted to 0.5.

**Table 1.** Value of the parameters of the controllers for step length and duration

| *Parameter*<br>*Controller* | $P$ | $D$ |
| --- | --- | --- |
| $C_{K_s}$ | -20 | 18 |
| $C_{pu_s}$ | 0.1 | 0.9 |

The model shows no hip retraction for walking on the flat floor at *ϕ*_0_ = 0 (Fig.14). Consequently, this model was chosen as the model with no hip retraction. As it is demonstrated in Fig.10 and Fig.12, the degree of SLR grew with increasing the magnitude of *ϕ*_0_. Fig.15 displays the range of stable walking on different ramps for each value of *ϕ*_0_.

**Fig. 14.**
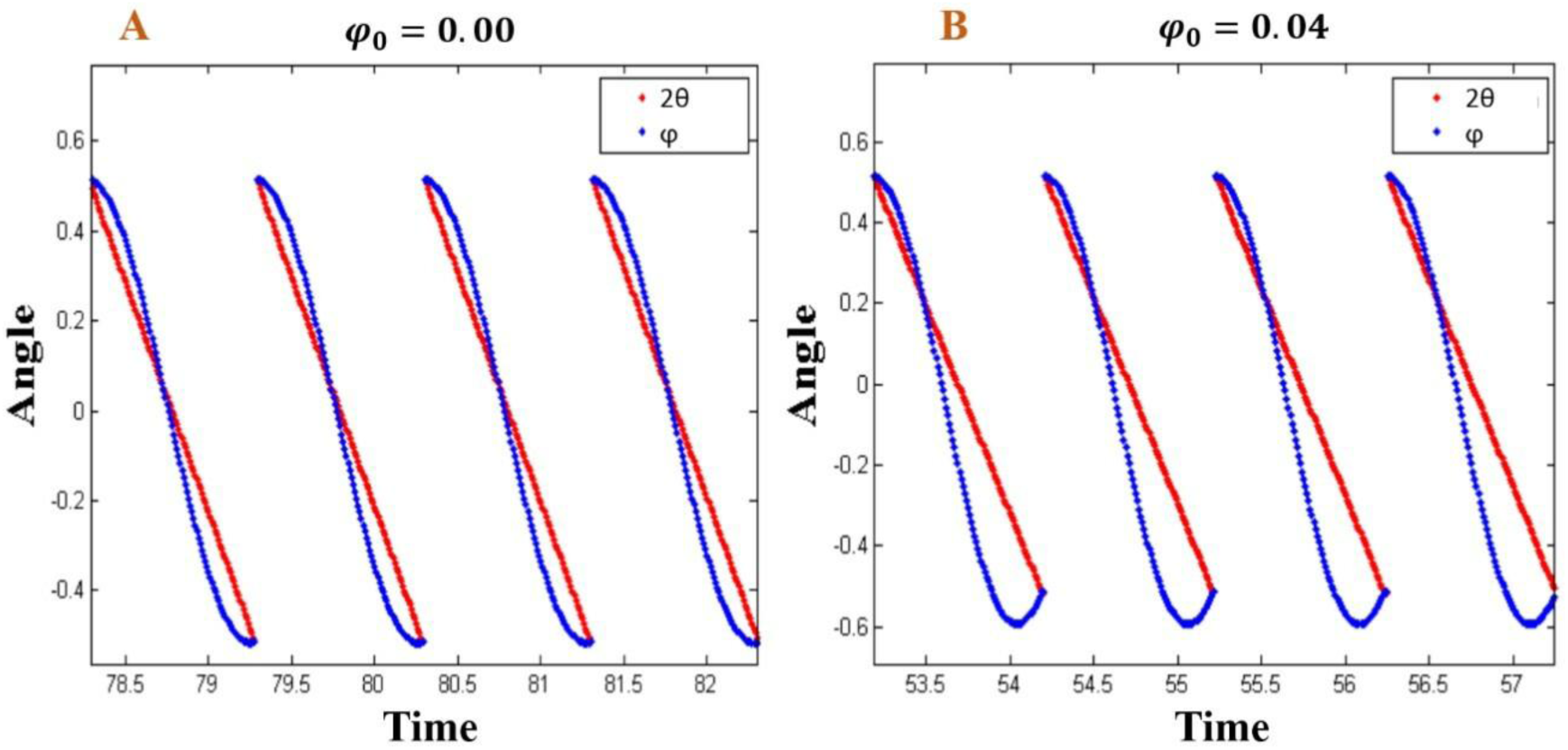
Swing and stance angles of the model under the influence of the controllers for the step length and step duration. To make a better illustration of the stopping condition of the single support phase (Eq.(4)), 2*θ* and *ϕ* were plotted in red and blue, respectively. A) *ϕ*_0_ = 0 and the model had no retraction, B) *ϕ*_0_ = 0.04 and the model retraction was seen obviously.

**Fig. 15.**
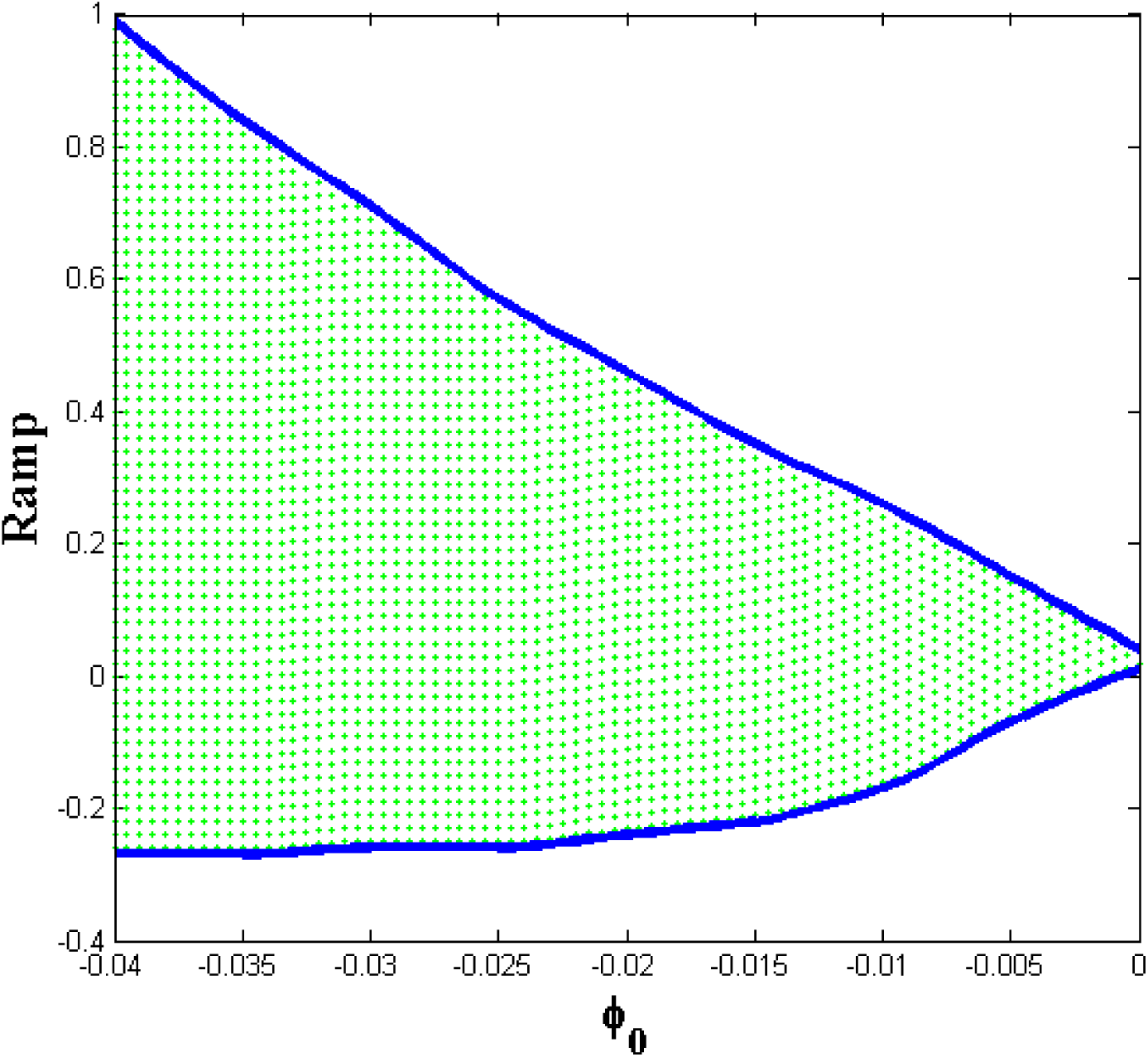
Range of the stable ramp angle for each *ϕ*_0_ is displayed by the green area.

## 4. Discussion

Hip retraction is a phenomenon that is seen in animals and humans during walking and running. The purpose of this study was to investigate the relationship between this phenomenon and the equilibrium point hypothesis, which is widely considered as a theory that explains how humans and animals control their body [36, 37]. In this paper, analyzing the experimental data revealed that the hip retraction increases during the walking on the ascending and descending stairs compared to the one during walking on flat floor. Thus, in this study, the effect of hip retraction on positive and negative ramps was investigated. Our proposed model shows that the hip retraction enhances the walking stability on the ascending and descending ramps.

In a stepping forward task, it was reported that a global EMG minima existed in the mid-swing when the swing hip was located just a little more forward than the stance hip [30]. In the present study, considering the referent of the swing hip to be a little ahead of the stance hip, it is shown that the hip retraction emerges as a natural response of the walking system without the use of direct motion or torque programming. Therefore, the results of our proposed model confirm that of the aforementioned research on experimental data of global EMG minima.

In this study, investigation of the experimental data revealed more hip retraction during the ascending and descending walks than during the walking on flat ground. So, it seems that the central nervous system initially increases the hip flexion angle to gain dominance over the periphery of foot placement, then hip retraction happens. The increase in the hip flexion angle and following hip retraction can be explained by a simple change on the position of the hip equilibrium point. However, some differences exist between the ascending or descending stair and ramp. Getting inspirations from the rise in the hip retraction during the ascending and descending walks on the stair, the stability of simple walking with and without hip retraction model for positive and negative ramps were compared. For the future work, it is recommended to investigate the effect of hip retraction on stepping on the stair for more complex models of a biped, which contain knee joint.

By setting certain values of *k* and *Impulse* for the model, certain step duration and length were achieved for walking on the flat ground. However, changing the ramp angle made alterations in the amount of step length and duration. In order to compare the walking stability on a ramp with and without hip retraction, controllers for step length and duration were designed. The controllers changed *k* and *Impulse* to make the step duration and length fixed. Fig.10 and Fig.12 show that the change of *Impulse* and *k* have meager effects on hip retraction. Consequently, changing them for controlling the step length and duration has negligible influences on the hip retraction.

Defining an equilibrium point angle for hip at just a little ahead of the stance leg results in an increase of hip retraction and rises the range of ramp angles for which the model can represent a stable walking. For instance, for *ϕ*_0_ = 0 and ramp angles within the range of [0.00, 0.04], the proposed model can simulate the walk and reach its proper step length and duration, while it is unstable out of this range. For *ϕ*_0_ = 0.04, the range is expanded to [−0.27, 0.99]. Therefore, expansion of the stability region may be assumed as a reason that the hip retraction during the walk on the ascending and descending ramps is more than that of the flat floor.

It has been frequently shown that the brain has a minimum role in the control of walking [38, 39]. Some animals, such as spinal cats, can walk after brain control command was completely disconnected from their limbs [40]. An advantage of the equilibrium point hypothesis over some other motion theories lies in its ability to explain how the brain controls the limbs with its minimum role. In this study, it is demonstrated that setting the equilibrium point at just a little ahead of the stance leg increases the hip flexion and then the hip retraction happens. Therefore, the maximum magnitude of hip flexion and hip retraction can be controlled by sending a simple control command.

## 5. Conclusion

In this study, it is demonstrated that hip retraction in humans during stride on ascending and descending stairs goes higher than that of the stride on a flat floor. Furthermore, the proposed model showed that during the walking, the proper definition of equilibrium for hip can result in a hip retraction emergence, increase, or decrease with no need to motion planning. It is also demonstrated that the hip retraction enhances the walking stability on a ramp. Thus, based on the equilibrium point hypothesis, it is shown that CNS needs to define equilibrium point just ahead of the stance leg to benefit from the hip retraction influence on ascending and descending walks on a ramp.

## Declarations of interest

none.

